# FiNN: A toolbox for neurophysiological network analysis

**DOI:** 10.1101/2022.02.11.479403

**Authors:** Maximilian Scherer, Tianlu Wang, Robert Guggenberger, Luka Milosevic, Alireza Gharabaghi

## Abstract

Recently, neuroscience has seen a shift from localist approaches to network-wide investigations of brain function. Neurophysiological signals across different spatial and temporal scales provide insight into neural communication. However, additional methodological considerations arise when investigating network-wide brain dynamics rather than local effects. Specifically, larger amounts of data, investigated across a higher dimensional space, are necessary.

Here, we present *FiNN* (*Fi*nd *N*europhysiological *N*etworks), a novel toolbox for the analysis of neurophysiological data with a focus on functional and effective connectivity. FiNN provides a wide range of data processing methods, and statistical and visualization tools to facilitate inspection of connectivity estimates and the resulting metrics of brain dynamics. The Python toolbox (https://github.com/neurophysiological-analysis/FiNN) and its documentation (https://neurophysiological-analysis.github.io/FiNN/) are freely available.

We evaluated FiNN against a number of established frameworks on both a conceptual and an implementation level. We found FiNN to require much less processing time and memory than other toolboxes. In addition, FiNN adheres to a design philosophy of easy access and modifiability, while providing efficient data processing implementations. Since the investigation of network-level neural dynamics is experiencing increasing interest, we place FiNN at the disposal of the neuroscientific community as open-source software.

## 1. Introduction

Analyzing dependence between neurophysiological signals, and the definition of large-scale networks, has become an important field of research that greatly enhances our comprehension of communication between distinct neural structures (Bressler & Menon, 2010; Siegel et al., 2012). Neural connectivity in particular is commonly quantified by estimating the degree to which neural oscillations within the same frequency band or across different frequency bands relate to each other (Fries, 2005). These two types of communication modes are known as same-frequency coupling and cross-frequency coupling, respectively (Friston, 2011; Hyafil et al., 2015).

Neural communication on a network level can be quantified on the basis of neurophysiological data from a wide variety of data sources including electroencephalography (EEG), magnetoencephalography (MEG), and local field potentials (LFPs) (Engel et al., 2013; Ganzetti & Mantini, 2013). An estimation of connectivity across regions and/or frequencies is generally more computationally intensive than the local synchronization of neural activity within a smaller spatial region (He et al., 2019). While the number of power estimates are linearly related to the number of sensors the number of potential connectivity candidates increases in a quadratic order of magnitude. Furthermore, in many applications, the amount of neurophysiological data is rising rapidly, partly due to increasingly dense sensor setups and high sampling rates (Brinkmann et al., 2009; Sahasrabuddhe et al., 2020; Song et al., 2015), as well as to more demanding analysis techniques, such as machine learning. A greater number of samples is therefore required (Glaser et al., 2019; Kus et al., 2004).

In recent years, a number of toolboxes include functions for estimating neural connectivity. However, the majority are either heavyweight frameworks that deeply encapsulate data, making modifications or the integration into an existing pipeline difficult and increasing the time required for memory processing; or frameworks with broad usability, but limited functionality for neuroscientific data analysis and interpretation (see Section 3). Here, we introduce FiNN (*Fi*nd *N*europhysiological *N*etworks), a novel framework written in Python that provides tools to analyze neurophysiological data in a bid to quantify network-wide neural communication within a lightweight and computationally efficient framework. FiNN includes several functions for cleaning and processing neurophysiological data in the context of connectivity. It also includes visualization routines and statistical methods, both of which are useful tools for the analysis of connectivity in large, high-dimensional neurophysiological data sets.

The goal of FiNN is to provide an open-source software toolbox offering easy-to-use and computationally efficient methods for both users and developers. From a user perspective, it is important that the toolbox is accessible and manageable. We therefore designed the functions in FiNN such that they can be readily used without a deep knowledge of the underlying functionality. In addition to an elaborate documentation on the individual functions, we included detailed information on the internal processing functions to promote modifiability. Furthermore, to facilitate the analysis of large datasets across high dimensional spaces, memory and CPU consumption were strictly monitored and reduced, thereby achieving a high level of scalability.

Section 2 describes the functionality of the toolbox, while Section 3 discusses FiNN in relation to the established toolboxes generally used to analyze neurophysiological data. This is followed by an illustration of its performance in comparison to a selection of established toolboxes in Sections 4 and 5.

## 2. Materials & Methods

### 2.1. Toolbox documentation and installation

FiNN (v1.0) is freely available to the research community as open-source code on GitHub (https://github.com/neurophysiological-analysis/FiNN). It can be downloaded as a zip file containing the last release, or by cloning the git repository. Documentation is available at https://neurophysiological-analysis.github.io/FiNN/, and includes a detailed description of the functions implemented. Furthermore, FiNN contains a demo folder with several trial scripts. The scripts are intended to provide an introductory demonstration of the functions implemented, but can also be used to gain a deeper methodological understanding of the functions. An exemplary workflow utilizing FiNN for the analysis of EEG data is provided in Table 1. A real-world application of FiNN can be observed in (Scherer, Steiner, et al., 2022) where FiNN was used to quantify phase-amplitude coupling between neuronal spiking and LFP activity. In the scope of this work, FiNN was used for data processing & statistical analysis.

**Table 1.**
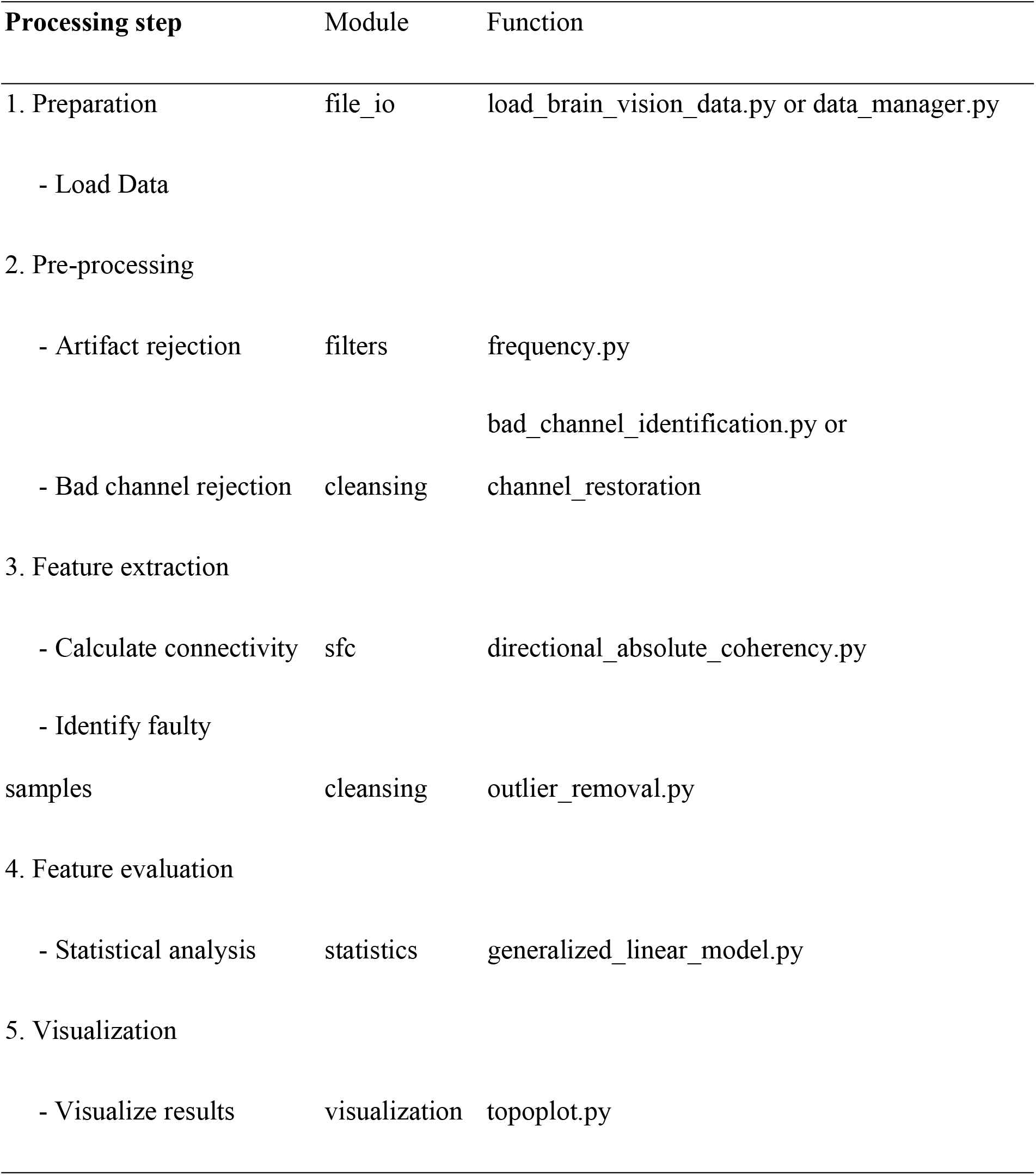
Illustrative example of a pipeline for processing raw neurophysiological data.

| Processing step | Module | Function |
| --- | --- | --- |
| 1. Preparation | file_io | load_brain_vision_data.py or data_manager.py |
| - Load Data |  |  |
| 2. Pre-processing |  |  |
| - Artifact rejection | filters | frequency.py |
|  |  | bad_channel_identification.py or |
| - Bad channel rejection | cleansing | channel_restoration |
| 3. Feature extraction |  |  |
| - Calculate connectivity | sfc | directional_absolute_coherency.py |
| - Identify faulty |  |  |
| samples | cleansing | outlier_removal.py |
| 4. Feature evaluation |  |  |
| - Statistical analysis | statistics | generalized_linear_model.py |
| 5. Visualization |  |  |
| - Visualize results | visualization | topoplot.py |

The FiNN toolbox was developed with Python version 3.8 (Van Rossum & Drake Jr, 1995) and requires the following Python packages: numpy (Harris et al., 2020), scipy (Virtanen et al., 2020), PyQt5, functools, lmfit (Newville et al., 2016), matplotlib (Hunter, 2007), rpy2. The MNE package (Gramfort et al., 2013) is required if the user wishes to load BrainProducts data (BrainProducts GmbH, Gilching, Germany). The following R (R Core Team, 2021) libraries are required if the user wishes to perform statistical evaluation: Matrix (Bates & Maechler, 2021), lme4 (Bates et al., 2015, p. 4), carData (Fox et al., 2020), and car (Fox & Weisberg, 2019).

### 2.2. Organisation of the FiNN Toolbox

FiNN (v1.0) consists of nine modules: *basic processing, artifact rejection, filters, cross-frequency coupling, same-frequency coupling, statistics, visualization, file IO*, and *miscellaneous*. A brief description of each module is provided below. Further details can be found in the online documentation (https://neurophysiological-analysis.github.io/FiNN/). Recommendations are also provided as to how to apply these for analysis, where applicable.

#### 2.2.1. Basic processing package

For initial processing, the basic processing package offers the *common average re-referencing (CAR)* and *downsampling* functions. This function subtracts from the data the part that is shared by all channels, since it is presumably the result of activity at the reference electrode, and hence equal in all channels. This procedure is recommended only when a large number of EEG channels (≥ 64) is available and these are equally distributed across the head (Nunez, 2010). The *downsampling* function downsamples a signal from its original sampling frequency to a lower, configurable target sampling frequency. Prior to the application of the *downsampling* function, it is important for the signal to be lowpass filtered below half the target frequency, as aliasing artifacts may otherwise be introduced into the downsampled signal (Shannon, 1948, 1949). Furthermore, we recommend that the target sampling frequency be set as low as possible while maintaining a sufficient level of temporal accuracy to accelerate data evaluation further down the line.

#### 2.2.2. Artifact rejection package

The artifact rejection package contains two functions to detect artifacts, and one function to clean the data. *Bad channel identification* identifies individual channels in which the power in a predefined frequency range deviates by more than three standard deviations (default value; configurable) from the mean power of all given channels as faulty channels, since spectral power is a useful feature for separating artifacts from EEG (Islam et al., 2016). In an *optional* second step, the function provides a custom-built dialogue window that shows the mean power of each channel (Figure 1A). Channels marked as faulty are automatically highlighted, and the user can visually confirm the selection and make changes accordingly. The outputs of the *bad channel identification* function are a list of non-faulty channels, and a list of faulty channels. Furthermore, it is highly recommended that frequency bands that are part of the subsequent analysis for artifact detection be avoided. This advice should be followed to ensure that the results are not biased, as side effects may arise if the power in the frequency band of interest is used in the artifact rejection.

**Figure 1.**
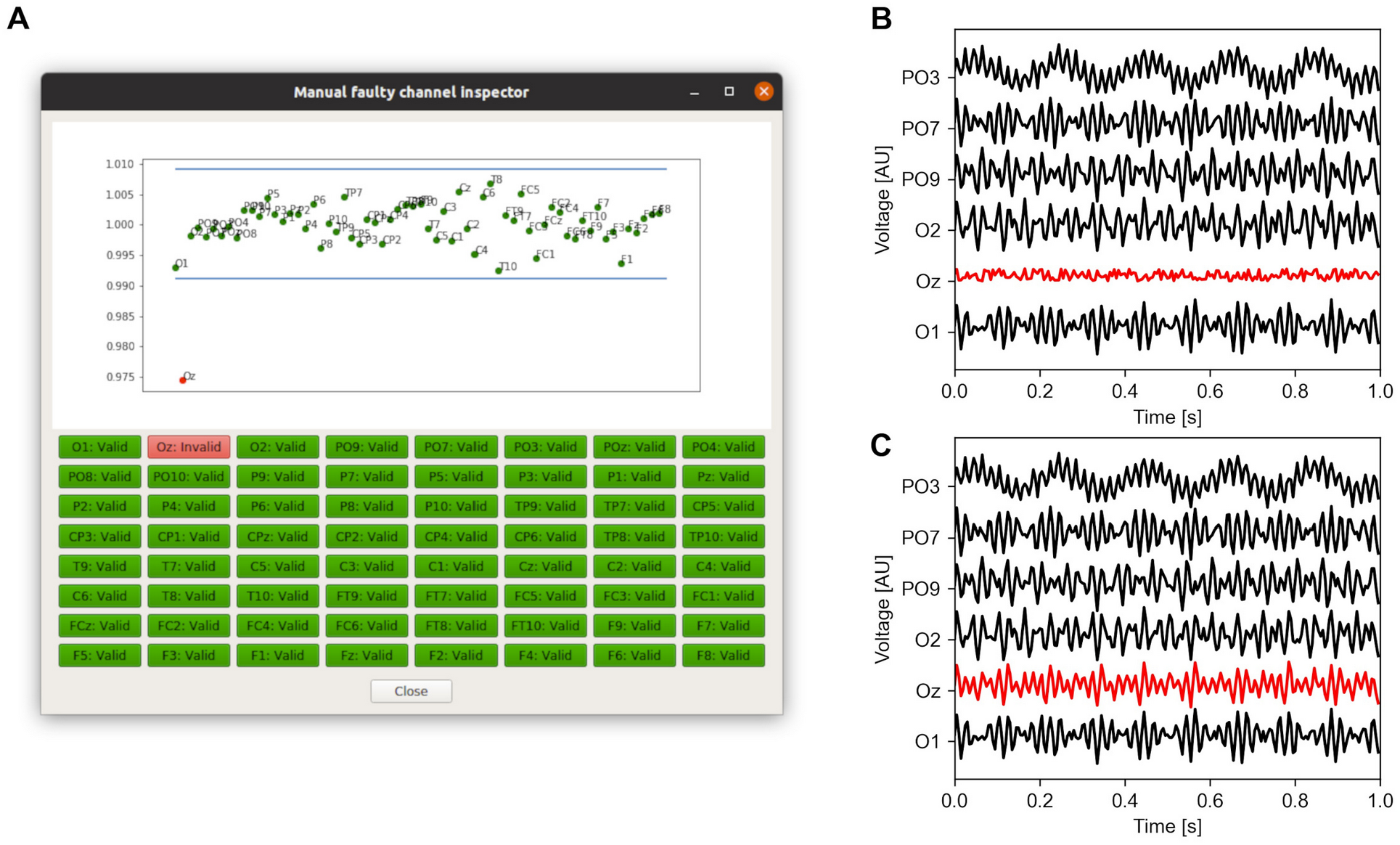
Practical application of the bad channel detection and channel restoration functions. (**A**) Bad channel detection. This figure shows the average power in a predefined frequency band for all included channels. A channel is flagged as bad if the individual power-based z-score is more than 3 standard deviations from the mean. Channels can be manually classified as normal or faulty either by clicking the corresponding dot in the plot, or by pressing the corresponding button. (**B**) Time-series traces at a sub-selection of the EEG electrodes shown in (A) before channel restoration. (**C**) Time-series traces at the same sub-selection of EEG electrodes shown in (B) after channel restoration.

Once the faulty channels have been identified, an optional follow-up step is to restore them using the *channel restoration* function. This restores faulty channels by averaging the signals from their respective neighbors (Figure 1B, 1C). A default adjacency matrix is provided to the *channel restoration* function. In the event of one or more neighboring channels of a faulty channel being faulty themselves, the channels are iteratively restored, commencing at the channel with the most non-faulty neighbors. Once a channel has been restored during an iteration, it is considered a non-faulty channel during the next iteration of the reconstruction process.

The *outlier removal* function removes any sample within a data set with a z-score higher than two (default value, configurable). This process is repeated iteratively until only those samples with a z-score of less than two remain or until a minimum sample threshold is reached. This function should be applied only to data from a unique, single state (e.g., within the same subject and same condition). It assumes that provided data is from a single process with a Gaussian distribution. Any data segments which fail this assertion are iteratively removed from the provided data. In the event that this assumption cannot be met, the approach presented here may not apply. The assumption of Gaussianity may be evaluated by either visual inspection or tests for normality.

#### 2.2.3. Filters package

The main function in this module is the implemented *finite impulse response (FIR)* filter. The filter is implemented via an overlap add approach (Rabiner & Gold, 1975) to speed up the processing procedure, especially for longer signals. Furthermore, the filter implementation provides a rapid and precise operation mode which initially converts the input data into either 32 bits floats (fast) or 64 bits floats (precise), and subsequently performs all operations with the required precision. The FIR filter can be configured as either a low-pass, high-pass, band-pass or band-stop filter. Furthermore, custom filters can be easily designed and accessed using the functions listed in the main FIR function. Additionally, a wraparound scipy’s Butterworth filter is also available.

#### 2.2.4. Cross-frequency coupling (cfc) package

The cross-frequency coupling package currently implements the following phase amplitude coupling (PAC) metrics: direct modulation index (Scherer, Wang, et al., 2022a), modulation index (Tort et al., 2008), phase-locking value (Mormann et al., 2005), and mean vector length (Canolty et al., 2006). A description of the metrics can be found in Table 2. The input signals should already be filtered into the frequency bands of interest, e.g., with the FIR filter implemented in the filters package (Section 2.2.3).

**Table 2.**
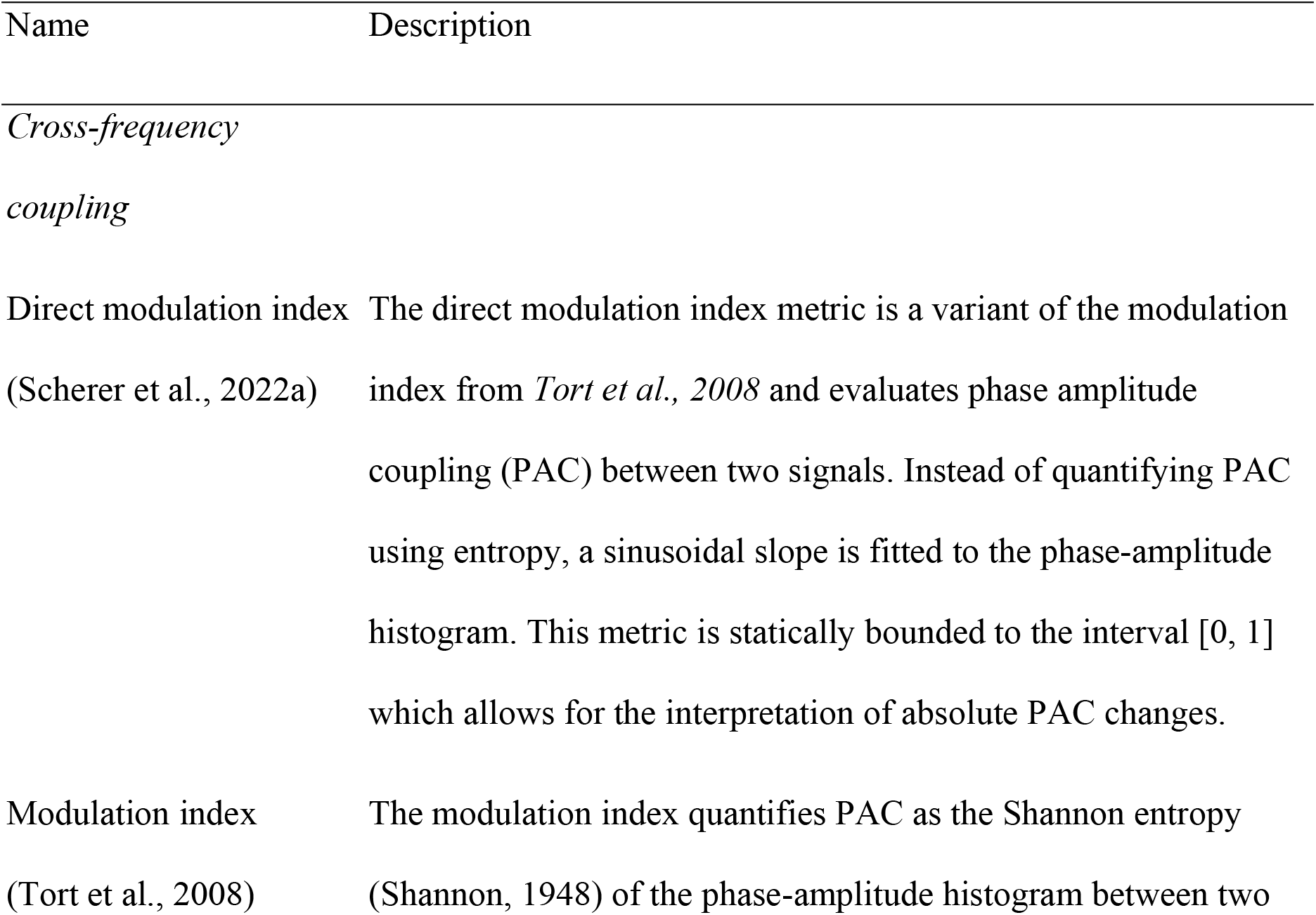
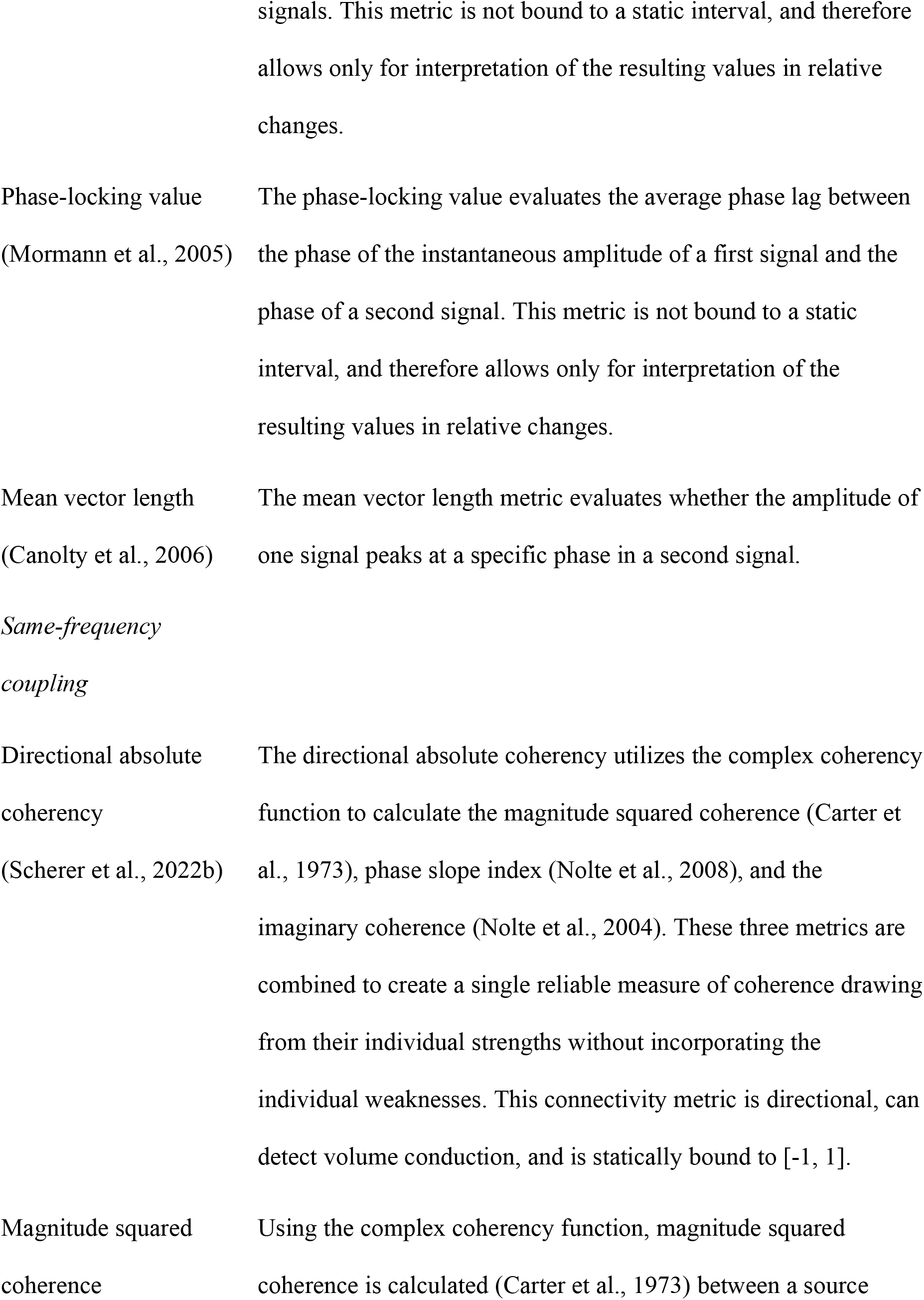
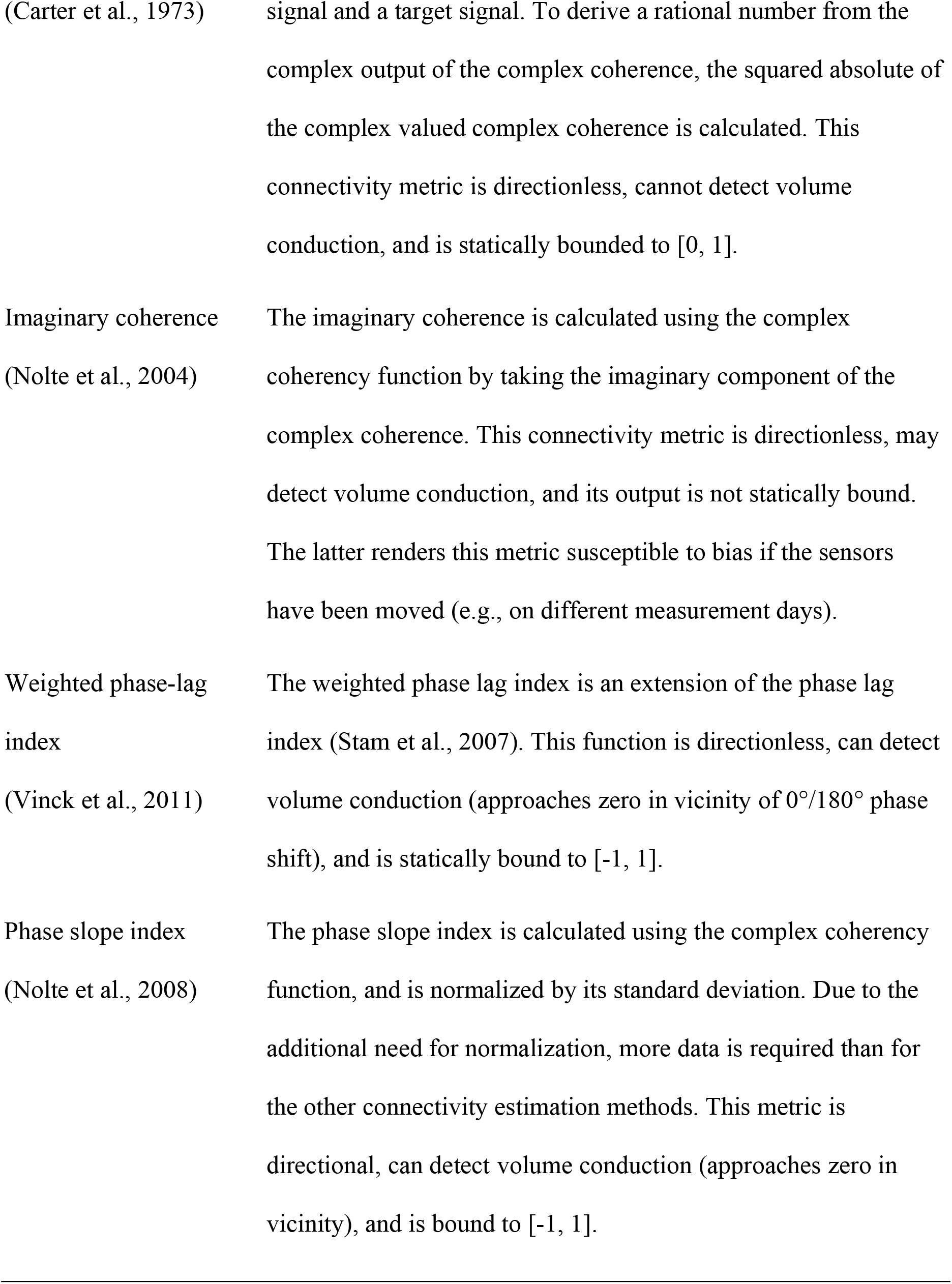
Overview of the cross-frequency and same-frequency coupling metrics implemented in the FiNN Toolbox.

#### 2.2.5. Same-frequency coupling (sfc) package

The same-frequency coupling package currently implements the following metrics: directionalised absolute coherency (Scherer, Wang, et al., 2022b), magnitude squared coherence (Carter et al., 1973), imaginary coherence (Nolte et al., 2004), weighted phase lag index (Vinck et al., 2011), and phase slope index (Nolte et al., 2008). A description of the metrics can be found in Table 2. It also includes a function for calculating the complex coherency, which can be interpreted as a measure of consistency between two signals with a constant phase shift, at a specific frequency (Shaw, 1984). Complex coherency is implemented as an additional function since it is a common precursor of the magnitude squared coherence, imaginary coherence, and others. The metrics implemented in the sfc module accept input data from the time domain, the frequency domain, and the complex coherency domain. Apart from the most commonly used time domain signals, the additional input data formats (frequency domain and complex coherence domain) were added to support modifiability of the implemented functionality, and to allow more efficient processing of multiple connectivity metrics in parallel when the data is already available in the required format.

#### 2.2.6. Statistics package

The statistics package contains the *generalized linear mixed models* function, which allows to employ generalized linear mixed models in the statistical evaluation of an investigation. The rpy2 package is used to wrap around the lme4 package (Bates et al., 2015) in *R (R Core Team, 2021)*. The implementation presented in FiNN provides a complete and easily interpretable output which comprises both the significance values, indicating how reliable an effect is, and the coefficients, allowing for an estimation of how impactful an observed effect is.

#### 2.2.7. Visualization package

FiNN offers the *topoplot* function, with the additional functionality of visualizing various levels of significance. Depending on their statistical significance, individual channels may be marked both before and/or after multiple comparison correction (Figure 2). The *topoplot* function is built on top of Matplotlib (Hunter, 2007), which is a data visualization library in Python. Since this function is solely responsible for visualization, any feature calculation and/or statistical evaluation has to be performed independently.

**Figure 2.**
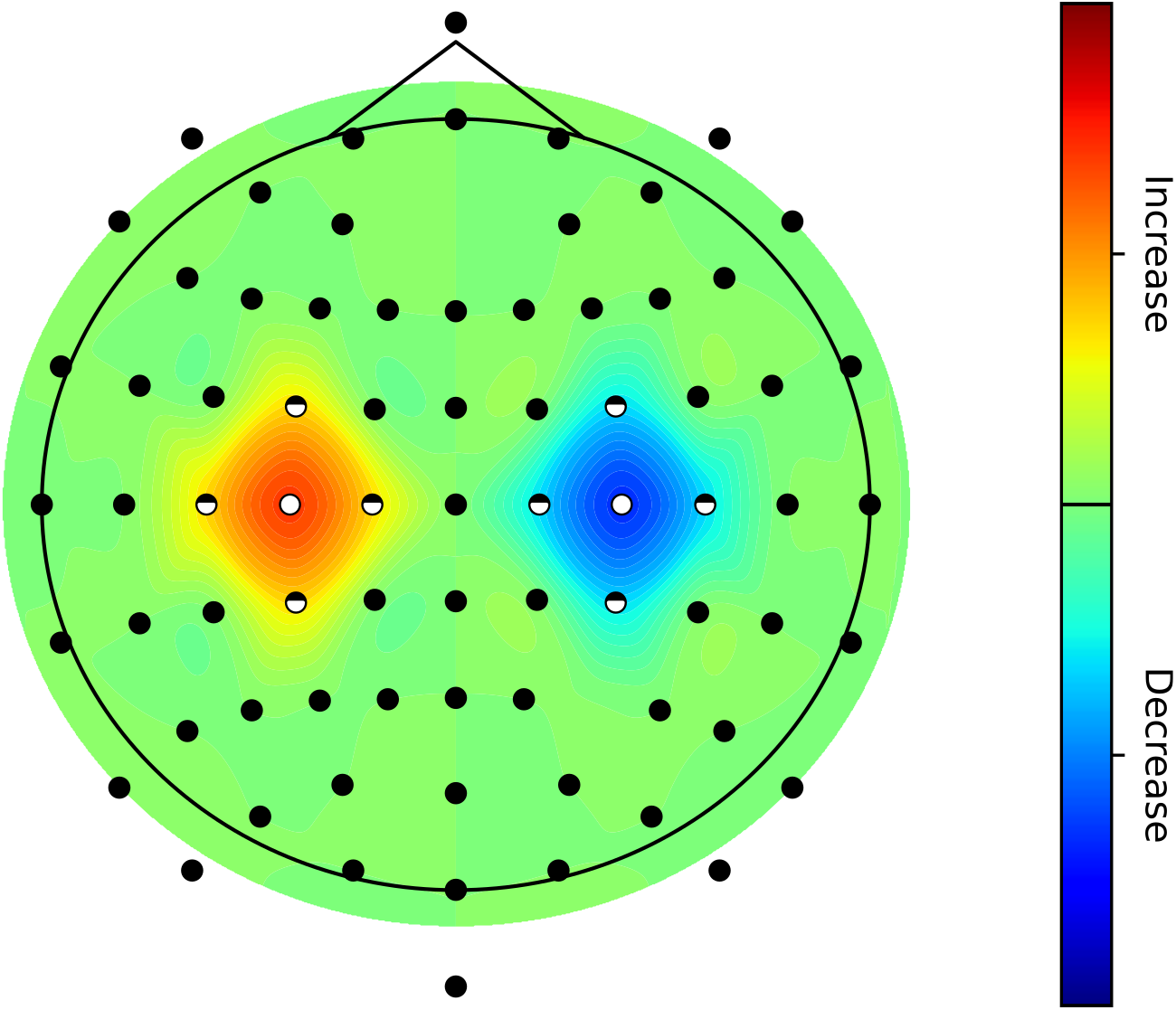
Topography plot of artificial demo data. This figure shows an example of a significantly increased activity above the left motor cortex and significantly decreased activity above the right motor cortex. At the center, the effects were modeled to be significant after multiple comparison correction. This is marked at the electrodes position using full white dots. The outlying areas of these effects were simulated to be also significant, but only prior to multiple comparison correction, this is indicated by half-filled dots at the electrodes location. Finally, non-significant areas are indicated by black dots.

#### 2.2.8. File IO package

When investigating network-wide brain dynamics, larger amounts of data have to be accommodated. FiNN offers the *data manager* module to handle large file sizes. In Python, pickle (.pkl) is a well-known tool for saving arbitrary variables to disk. However, when loading or writing a file, it has the well-known shortcoming of consuming multiple times more memory than the size of the file itself. This causes memory spikes which may in turn lead to unstable behavior and crashes if not handled carefully. The *data manager* function circumvents this issue by processing a large file into several smaller ones. This increases the stability of the overall processing pipeline and reduces the likelihood of requiring user intervention, particularly if the pipeline is partially or fully automated.

#### 2.2.9. Miscellaneous package

Due to the large size of neuroscientific data sets, it is beneficial to separate the processing into subprocesses and perform operations in parallel. When using the Python native subprocess pool, however, all subprocesses may receive or send their data at the same time. This behavior is similar to issues of pickle-based file reading and writing (see previous section). The FiNN Toolbox therefore offers the *timed pool* function, a custom-built subprocess pool that limits the sending or receiving of data to one subprocess at a time. This implementation substantially decreases the risk of memory spikes for a negligible increase in run time. Additionally, the *timed pool* function has two advantages over the Python native subprocess pool. First, there is an option to delete the original input data immediately after its function is executed, thus releasing additional memory. Second, there is an option to add a configurable life-time duration to each subprocess. When a subprocess does not return a result within the allotted life-time duration, it will be restarted. This behavior is particularly suitable in the event that one or more of the subprocesses do not terminate within a reasonable time, e.g., when running a randomly initialized optimization loop.

### 2.3. Code testing and validation

System tests have been successfully applied to all the functions implemented in FiNN. Functional tests were performed for mathematically more complex components such as the FIR filter and the complex coherence calculation using scipy as a reference implementation. Automated unit tests were implemented in the framework to verify that its behaviour is as designated.

### 2.4. Performance evaluation

We evaluated the performance of the FIR filter against the implementation from MNE (a Python based framework for the analysis of electrophysiological data, fully defined in section 3.1) in terms of speed, and the performance of the subprocess pool against both the implementation from *joblib* used in MNE and the default subprocess pool implemented in the multiprocessing package in Python, in terms of RAM consumption. These two functions were selected as they necessitate efficient implementations (both RAM and CPU time-wise) if evaluation is to be rapid. Biological data recorded from the human subthalamic nucleus was used for the evaluation process (Milosevic et al., 2020). During the three subprocesses, signals of different lengths, varying between 30 seconds and 24 hours (artificially expanded), were filtered using both FIR implementations. The data was appended via repetition as required. All parameters were configured equally to achieve a maximum degree of comparability between the two implementations. The scripts of the above evaluations and comparisons are provided in the toolbox (https://github.com/neurophysiological-analysis/FiNN).

## 3. Results

### 3.1. Comparison to other frameworks

A large number of open-source toolboxes have been developed to support the processing and analysis of (neurophysiological) signals. Here, we compare FiNN to a selection of existing toolboxes in terms of scope and computational performance (i.e., processing time and memory consumption). These other toolboxes were selected either on account of their reputation in scientific data analysis (e.g., scipy) or as a result of key-word-based searches (e.g., “Python toolbox EEG”). We selected the search engine *startpage*.*com*, as it delivers Google results while anonymizing search requests, and is therefore not subject to a user-specific filter bubble (Salehi et al., 2018). In the following paragraphs, the selection of frameworks with which the FiNN framework is compared is described in more detail.

Two Python-based frameworks that are widely used in general data processing and analysis are *scipy (scientific python) (Virtanen et al*., *2020)* and *scikit-learn (Pedregosa et al*., *2011)*. Scipy and scikit-learn are focused on fundamental algorithms (e.g. optimization techniques, basics in digital signal processing, and linear algebra), and machine learning/data analysis, respectively. Currently, scikit-learn provides functionality such as the principal component analysis (Hotelling, 1933; Pearson, 1901) and the independent component analysis (Makeig et al., 1995), which are commonly used for the purpose of artifact identification and removal (Xue et al., 2006).

A major advantage of frameworks such as scipy and scikit-learn is that they are usually heavily modified towards a small memory footprint and limited CPU consumption. Scikit-learn in particular has been optimized for speed, which becomes increasingly important as the complexity of the applied machine learning algorithm increases. Relative to the other frameworks discussed in this paper, scipy may be categorized as a low-level framework. Generally, scipy and scikit-learn offer a great range of functions which can be – and often have to be – highly customized to a problem at hand. This in turn requires in depth knowledge of the provided methodology in the respective areas. Furthermore, while scipy and scikit-learn excel with regards to the provided functionality (fundamental algorithms and machine-learning/data analysis), on their own, these frameworks do not fully cover all needs of electrophysiological data analysis. For example, scipy/scikit-learn lacks some data analysis functions elementary in many experimental, electrophysiology based neuroscientific analyses, such as connectivity metrics, and proper visualization functions, such as topoplots. Having a different focus, FiNN uses the comparatively low-level functionality provided by scipy (and other toolkits) to provide this functionality for EEG/MEG data analysis.

Another broad Python-based framework specifically developed for the analysis and visualization of neurophysiological data, is *MNE (Gramfort et al*., *2013)*. MNE has its roots in the estimation of source-space signals from signal-space signals via EEG or MEG recordings. Unlike scipy and scikit-learn, MNE is a very high-level framework that offers a wide range of functions to analyze neurophysiological data. These methods are often configured in predefined ways, making any deviations from the intended application case rather difficult. MNE focuses strongly on mathematical precision. This, in turn, mandates a high memory consumption and slow data processing speed. MNE follows a fundamentally different design philosophy to FiNN. While FiNN aims to provide methods which can be easily integrated into any workflow, MNE is almost exclusively used if the proposed MNE-specific pipelines are implemented for data processing. This results in a lower degree of customizability when using MNE rather than FiNN. Another difference is the focus of optimization. MNE focuses on optimal mathematical accuracy, whereas FiNN allows the user to define the trade-off between high mathematical accuracy, high processing speed and efficient resource usage, e.g., in applications where online analysis is necessary.

*NeuroKit2 (Makowski, 2016)* is another Python-based framework for the analysis of neuroscientific and neurophysiological data. The framework was designed to work with electrocardiography (ECG), electromyography (EMG), and EEG data. It provides an adequate range of functionality with a different focus to FiNN. While NeuroKit2 aims to be a generalist for any kind of neurophysiological signals, FiNN focuses on EEG/EMG/MEG and LFP signals. This difference in focus can be readily observed in the functionality provided. For instance, NeuroKit2 offers methods to analyze heart rate variability via ECG, or the autocorrelation of EEG signals, while FiNN offers metrics for connectivity between EEG channels, or functionality to visualize a topoplot. Neither connectivity nor topoplot functionality are offered in NeuroKit2.

Finally, there is a wide range of Python-based frameworks with limited functionality or rather rigid data processing pipelines. These frameworks include NeuroPycon (Meunier et al., 2020), Plotly (Plotly Technologies Inc., 2015), matplotlib (Hunter, 2007), HEAR (Kobler et al., 2019), Pygpc (Weise et al., 2020), Human Neocortical Neurosolver (Neymotin et al., 2020), Neo (Marcus et al., 2019), nipype (Gorgolewski et al., 2011), ScoT (Billinger et al., 2014), PyEEG (Bao et al., 2011), Gumpy (Tayeb et al., 2018). Due to a too limited overlap with the focus of FiNN, these frameworks were not compared to FiNN.

### 3.2. Performance comparison between FiNN and MNE

The FIR filter and the multiprocessing pool of FiNN were compared to the same functions when implemented in MNE. The processing times of the FIR filter as implemented in FiNN and MNE are shown in Figure 3. Both configurations in FiNN were substantially faster than their counterparts in MNE, provided that continuous data sets were of 1-hour length or less. When the fast implementation was chosen in FiNN, this implementation of the FIR filter was always executed more quickly than its MNE counterpart. While the difference was up to 670% more rapid for small data sets, the implementation of FiNN remained approximately 10% faster for very long continuous data sets (24 hours) (Figure 3). Performance evaluations were executed using Python v3.9 on a desktop PC (CPU: AMD Ryzen 9 5950x, RAM: 64 GB (2×32 GB DDR4), Mainboard: B550 AORUS ELITE AX V2).

**Figure 3.**
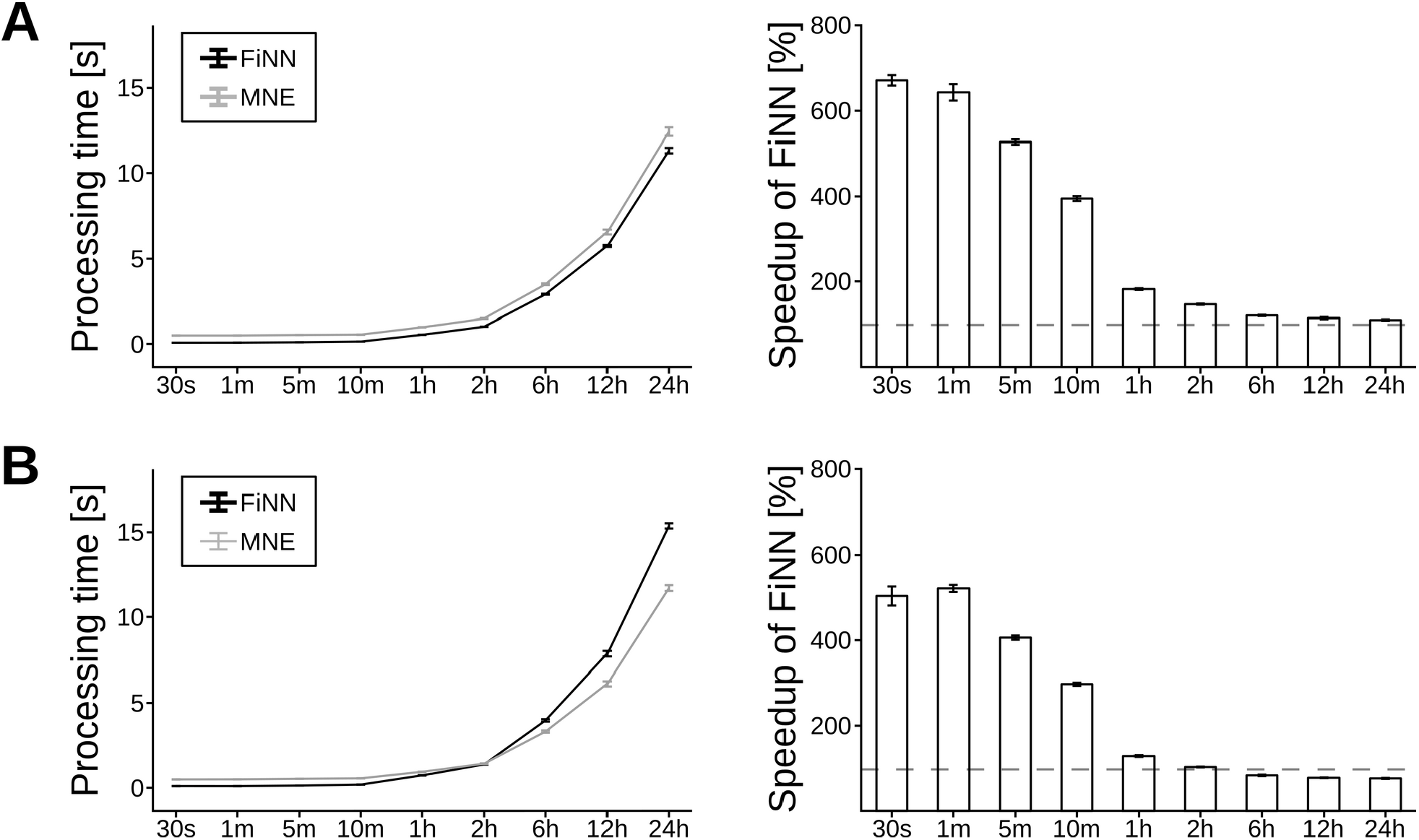
Processing time of the FIR filter implemented in FiNN and MNE. The rows show the processing time (mean and standard deviation) on signals with varying durations in the (**A**) fast mode and (**B**) precise mode. The left column shows the runtime in microseconds. The right column shows the percentage increase in speed of the FIR filter implemented in FiNN relative to the implementation in MNE (mean and standard deviation).

As shown in Figure 4, the multiprocessing pool implemented in FiNN requires less RAM compared to the default multiprocessing package included in the native Python multiprocessing package and MNE with the multiprocessing package backend.

**Figure 4.**
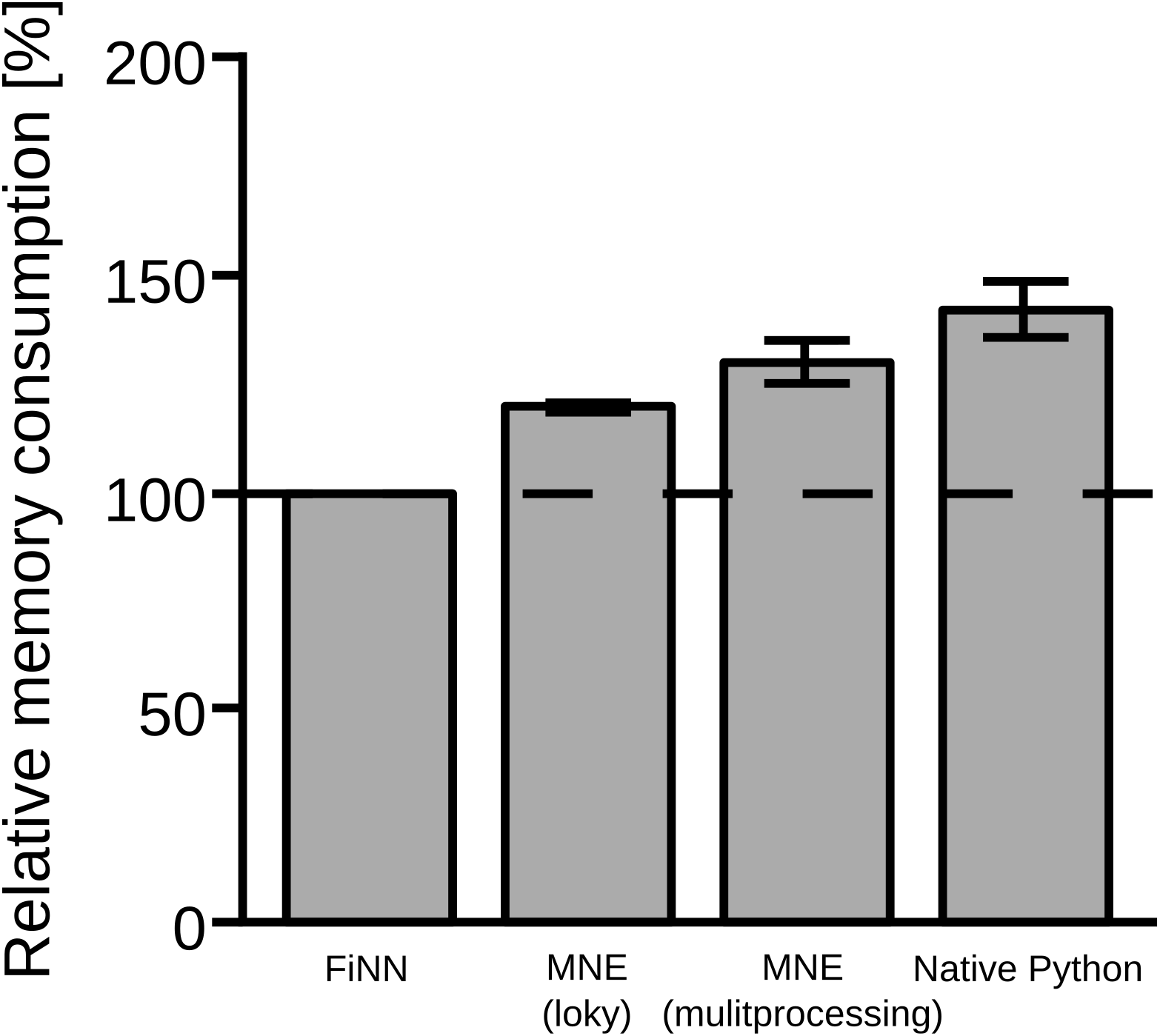
Random access memory of the subprocess pool as implemented in FiNN, MNE, and the native multiprocessing package in Python. For comparison, the relative memory consumption of each pool is shown as a percentage of the memory consumption of the multiprocessing pool implemented in FiNN.

## 4. Discussion

Neuronal information processing takes place on a local level and on a network level (Reimers, 2011; Thorpe, 1989; Zhang et al., 2016). Therefore, to understand how information is processed, it is also crucial to investigate the non-local components of information processing. To this end, we present FiNN, a Python based framework used to *Fi*nd *N*europhysiological *N*etworks in neurophysiological data and subsequently analyze them. FiNN implements both established and novel metrics for the evaluation of the same frequency coupling (Fries, 2005) and cross frequency coupling (Canolty & Knight, 2010) analyses used to investigate network level information flows. In particular, FiNN offers implementations for the newly proposed connectivity metrics directional absolute coherence (Scherer, Wang, et al., 2022b) and direct modulation index (Scherer, Wang, et al., 2022a).

The amount of data collection in experimental neuroscientific applications is steadily rising with increasingly powerful frequency amplifiers and an ever-growing number of simultaneously recorded channels (Song et al., 2015). Concurrently, the number of features extracted from this raw data also increases (Vaid et al., 2015). For these reasons, efficient data processing is essential. One major benefit of FiNN is its strong optimization towards processing speed and minimal memory consumption. This is reflected not only in the individual functions, which have been optimized to perform as little recalculation as possible, but also in the modules provided. Exemplary modules for this design philosophy are the custom subprocess pool for parallel processing or the data manager for I/O operations, both of which are included in this framework.

Despite the fact that FiNN includes many elements to assist potential users in the analysis of neurophysiological data, modifiability was still a major concern during development. Although the functionality available offers many parameters to calibrate it to a specific application case, edge cases are difficult to identify. The high level of modifiability provided by FiNN should be most helpful in these cases. Since all functionality is implemented in open-source languages, the programming code can be easily amended to cover specific cases encountered in a user’s data analysis. Furthermore, not only the uppermost, but any layer of functionality was fully documented to increase support for potential customization efforts.

Although the main focus of FiNN is on the analysis of network level communication (both same-frequency and cross-frequency) from neurophysiological data, it also provides a full evaluation pipeline for the investigation of local and network level information processing from EEG, MEG, EMG, and/or LFP based data. The pipeline offered by FiNN includes modules for data pre-processing such as semi/fully-automated outlier detection, postprocessing and visualization, and structural functionality to ease parallel processing. Although FiNN was originally developed with EEG and EMG data in mind, the implemented methods are well suited to analyze any kind of neurophysiological signals (e.g., LFP and MEG).

### 4.1. Outlook

Initially, FiNN was developed for in-house evaluation of experimental paradigms generating neurophysiological data. As the number of paradigms and their subsequent analyses increased, so too did the functionality of FiNN. Meanwhile, FiNN has been developed to the extent where it not only supports in-house evaluation of neuroscientific investigation, but also external ones. We are therefore pleased to share FiNN as an open-source software with the neuroscientific community.

## Abbreviations

cfc: cross-frequency coupling
ECG: electrocardiography
EEG: electroencephalography
EMG: electromyography
FIR: finite impulse response
LFP: local field potential
MEG: magnetoencephalography
PAC: phase-amplitude coupling
sfc: same-frequency coupling.

## Credit authorship contribution statement

**Maximilian Scherer**: Conceptualization, Methodology, Software, Writing – original draft, Writing – review & editing.

**Tianlu Wang**: Writing – original draft, Writing – review & editing.

**Robert Guggenberger**: Conceptualization, Methodology, Writing – review & editing.

**Luka Milosevic**: Conceptualization, Methodology, Writing – review & editing.

**Alireza Gharabaghi**: Conceptualization, Writing – review & editing, Funding acquisition.

## Acknowledgements

This work was supported by the German Federal Ministry of Education and Research [BMBF]. We acknowledge support by the Open Access Publishing Fund of the University of Tübingen.

## Code and data availability

The data and code supporting the findings of this study are available on https://github.com/neurophysiological-analysis/FiNN.

